# New fossil wasp species from the earliest Eocene Fur Formation has its closest relatives in late Eocene ambers (Hymenoptera, Ichneumonidae, Pherhombinae)

**DOI:** 10.1101/2021.11.22.469510

**Authors:** Noah Meier, Anina Wacker, Seraina Klopfstein

## Abstract

Darwin wasps (Ichneumonidae) are one of the most species-rich insect families, but also one of the most understudied ones, both in terms of their extant and extinct diversity. We here use morphometrics of wing veins and an integrative Bayesian analysis to place a new rock fossil species from the Danish Fur Formation (∼54 Ma) in the tree of Darwin wasps. The new species, *Pherhombus parvulus* n. sp., is placed firmly in Pherhombinae, an extinct subfamily so far only known from Baltic and Rovno-Ukranian ambers, which are estimated to be 34–48 Ma and 34–38 Ma, respectively. Our phylogenetic analysis recovers a subfamily clade within the higher Ophioniformes formed by Pherhombinae, Townesitinae and Hybrizontinae, in accordance with previous suggestions. Due to the placement of the new species as sister to the remaining members of Pherhombinae, we argue that our finding is not at odds with a much younger, late Eocene age (∼34–41 Ma) of Baltic amber and instead demonstrates that *Pherhombus* existed over a much longer period than previously thought. Our results also exemplify the power of wing vein morphometrics and integrative phylogenetic analyses in resolving the placement even of poorly preserved fossil specimens.

---

Insect taxonomy in the past centuries was strongly biased towards large and colourful species and thus overrepresented Lepidoptera and Coleoptera. More recently, other orders came into focus, especially Diptera and Hymenoptera, due to their extraordinary diversity and ecological and economic importance (Forbes et al. 2018, Ronquist et al. 2020). Among them, Darwin wasps (Ichneumonidae) are assumed to have one of the largest gaps between the number of described species and the actual species diversity (Klopfstein et al. 2019b). The fossil record of ichneumonids goes back to the Lower Cretaceous, about 120-130 Ma, while a recent dating study placed the origin of the family and most of its subfamilies in the Jurassic (about 181 Ma; Spasojevic et al. 2021). However, the fossil record of Darwin wasps is even more under-researched than their extant diversity, which impedes inferences about their past diversity and evolutionary history. In this study, we describe an approximately 54 Ma old ichneumonid rock fossil species from the Danish Fur Formation (Rust 1998). Its forewing venation with a large, rhombic areolet is rather rare among members of the family, both extant and extinct, and makes it unique among the known Fur Formation ichneumonids.

## Darwin Wasp Fossils from the Early Eocene Fur Formation

The Fur Formation is located in northwestern Jutland in Denmark, with its center on the islands of Fur and Mors. The 60 m thick sediments consists of porous diatoms and contains approximately 200 volcanic ash layers that were deposited right after the Paleocene-Eocene Thermal Maximum, about 54 Ma (Chambers et al. 2003, Westerhold et al. 2009). It is one of the oldest Cenozoic deposits of fossil insects in Europe (Larsson 1975) and interestingly, only winged insect forms have been found so far (Rust 1998). This is probably due to the distance of 100 km from the Scandinavian coastal line at the time of deposition. The recovered insects were either blown onto the open sea by storms (Larsson 1975) or showed long-distance migratory behavior (Ansorge 1993, Rust 2000). Rust (1998) mentioned two forms of Darwin wasps that were common among Fur insects: one dark, strongly sclerotized and one light, less sclerotized form. However, more recent work showed that these forms each included multiple species (Klopfstein in press). Currently, there are ten Darwin wasp species known from Fur, all of which are classified in the extant subfamily Pimplinae (Henriksen 1922, Klopfstein in press). So far, no species from any of the other 41 extant and five extinct subfamilies have been recorded from Fur, even though preliminary analyses indicate a much higher diversity (own observations). Given that Darwin wasps have recently been estimated to date back to the Jurassic (∼181 Ma) and most extant subfamilies have probably started diversifying by the Early Cretaceous (>100.5 Ma; Spasojevic et al. 2021), a much higher diversity would also be expected for the early Cenozoic.

## Candidate subfamilies: Mesochorinae and Pherhombinae

The forewing venation of the fossil in question, especially the rhombic areolet, suggests that the new fossil species belongs to either the extant Mesochorinae or the extinct Pherhombinae (Broad et al. 2018). With 863 extant and 8 fossil species (Yu et al. 2016), Mesochorinae are quite a large subfamily. Typical features include a straight, needle-like ovipositor, in most cases a large rhombic areolet, a deep glymma in the first metasomal segment, and extended, rod-like parameres in the male. As far as we know, Mesochorinae are obligate hyperparasitoids, using mostly Ichneumonidae and Braconidae larvae as primary hosts (Broad et al. 2018). Brues (1910) described eight fossil Mesochorinae species from the Florissant Formation, approximately 34 Ma (McIntosh et al. 1992) and one species from Baltic amber (Brues 1923). This latter species was later transferred to Pherhombinae, an extinct subfamily described more recently (Kasparyan 1988).

The monotypic Pherhombinae was established based on two species from Baltic amber, *Pherhombus antennalis* Kasparyan, 1988 and *P. brischkei* (Brues, 1923). In 2005, Tolkanitz et al. described *P. dolini*, the first Pherhombinae found in Ukrainian Rovno amber (Tolkanitz et al. 2005). And recently, Manukyan (2019) described three further *Pherhombus* species from Baltic amber, increasing the number of species in the subfamily to six. Kasparyan (1988, 1994) proposed a close relationship of Pherhombinae with the extinct Townesitinae and the extant Hybrizontinae and cited several character states as potential synapomorphies for this clade. In a recent phylogenetic analysis that included one species of Pherhombinae (*P. antennalis*), this grouping was indeed recovered among other subfamilies of the Ophioniformes group, although with a very sparse taxon sampling (Spasojevic et al. 2021). Interestingly, Manukyan (2019) suggested a crepuscular or nocturnal activity for the subfamily based on the somewhat enlarged, raised ocelli. As all extant subfamilies that include nocturnal species (Ophioninae, Mesochorinae, Tryphoninae and Ctenopelmatinae) belong to Ophioniformes, the placement of Pherhombinae in this group appears plausible. Regarding the biology of Pherhombinae, only little is known otherwise, although their short ovipositor might indicate that they attack exposed hosts, for instance larvae of Lepidoptera or Symphyta (Belshaw et al. 2003).

## Amber fossils and their controversial age

All Pherhombinae described so far were found as inclusions in Baltic and Rovno amber (Manukyan 2019). Age estimates of Baltic amber vary considerably (about 56.0 to 33.9 Ma: Ritzkowski 1997, Perkovsky et al. 2007, Bukejs et al. 2019); they are based on biostratigraphic analyses (pollen, spores, phytoplankton), lithographic analyses of surrounding sediment, and K-Ar age estimation of glauconites in the layers Blue Earth, lower Blue Earth and lower Gestreifter Sand (Weitschat and Wichard 2010, Sadowski et al. 2017). The uncertainty range is due to a controversy over whether Baltic amber is autochthonously deposited in upper Eocene layers (Standke 1998, Sadowski et al. 2017), or redeposited there while originating from the Lower or Middle Eocene (Schulz 1999, Weitschat and Wichard 2010). A recent study even suggests that Baltic amber was deposited in a periodic fashion between 45–35 Ma due to the transgression and regression of the sea into the amber-producing forests (Bukejs et al. 2019). Similarly controversial discussions are ongoing for the somewhat more precise age estimates of Rovno amber (37.8–33.9 Ma), with a trend in recent studies towards 37–35 Ma (Dunlop et al. 2019). Even though there is a possible overlap in the age estimates of Baltic and Rovno amber, Perkovsky et al. (2007) describe pronounced differences in their insect assemblages, which can be explained either by a different age, by the location on different land masses, or by different regional climatic conditions. The finding of a Pherhombinae rock fossil in the Fur Formation (earliest Eocene) could contribute as another piece of the puzzle to the discussion about the likely age of Baltic and Rovno ambers.

## A combined approach to fossil placement

To obtain a robust placement of the new fossil species among ichneumonid subfamilies, we combined morphometrics of the wing veins (Li et al. 2019) with a Bayesian phylogenetic analysis based on a dataset using both morphological and molecular data (Spasojevic et al. 2021). We also aimed to test Kasparyan’s (1988, 1994) hypothesis about a close relationship of Pherhombinae with Townesitinae and Hybrizontinae using an extensive taxon sampling and the addition of relevant morphological characters to a combined molecular and morphological matrix. In the light of our results, we describe the new fossil species in the genus *Pherhombus* and discuss the implications of this finding on the potential age of Baltic amber and on the quality of fossil placements based on combined Bayesian analyses.

## Materials and Methods

### Morphological study of Fur fossil

The studied rock fossil (FUR #10652) was found by Jan Verkleij in Ejerslev (Denmark) and is deposited at the Fur Museum in Nederby. Both part and counterpart were available and about equally informative. So far, no other specimens of this fossil species are known. Images of the dry fossil and of the fossil covered in 85% ethanol were made with the digital microscope *Keyence VHX 6000* at 200x magnification. Both stitching and stacking techniques were applied to enhance image quality. The interpretative line drawing was made with the open-source software GIMP. The drawing is based on both part and counterpart. Solid lines imply a higher certainty for the interpretation than dotted lines. Differences in line width are used to visualise larger and smaller structures and do not imply varying certainty.

Morphological terminology follows Broad et al. (2018), while abscissae of wing veins are denoted as in Spasojevic et al. (2018). The colour description is based on the colours visible in the fossil. The original colours of the species may differ from that.

### Morphometric analysis of wing venation

After measuring several linear measurements that had been used in earlier studies of ichneumonid morphology (Bennett et al. 2019, Klopfstein and Spasojevic 2019), we chose the two most promising ratios to distinguish Mesochorinae and Pherhombinae based on visual inspection. For Mesochorinae, we obtained measurements from eight species based on drawings from Townes (1971) and for Pherhombinae, we used three wing photographs from Manukyan (2019), in combination with direct examination of two of the species. The new fossil species was measured from the obtained photographs, using average values from both forewings. To obtain an even denser taxon sampling, we in a second step also included incomplete fossils, for which only forewings could be measured, namely six fossil Mesochorinae (Brues 1910) and the three additional Pherhombinae species (Tolkanitz et al. 2005, Manukyan 2019). For a complete list of taxa sampled and the data used from each, consider Supplementary File S1. Wing vein lengths were measured with ImageJ version 2.1.0 and a scatterplot was obtained in R (R Core Team 2014).

### Morphological and molecular matrix

To test alternative subfamily placements, we performed a Bayesian phylogenetic analysis based on a combined morphological and molecular matrix (Spasojevic et al. 2021) compiled for total-evidence dating (Pyron 2011, Ronquist et al. 2012a). To simplify and thus speed up the analysis, we only included one or two taxa from 31 of the 45 ichneumonid subfamilies (Supplementary File S1). For the two focal subfamilies, Mesochorinae and Pherhombinae, we increased the taxon sampling by newly coding the morphological characters and, for the former, complemented the dataset with sequence data for the genes 28S and CO1 from Genbank (Table 1). In Mesochorinae, we coded one or two species in each extant genus, while we added all six described Pherhombinae species and a hitherto undescribed species from Baltic amber. To test the hypothesis that Pherhombinae are most closely related to Hybrizontinae and the extinct Townesitinae, we sampled additional species from these two subfamilies (Table 1). We did not include the fossil Mesochorinae from Florissant Formation (Brues 1910) in the phylogenetic analysis, as their descriptions did not allow sufficient coding of morphological characters. However, we did include them in the morphometric analysis of the fore wing (see below).

**Table 1.**
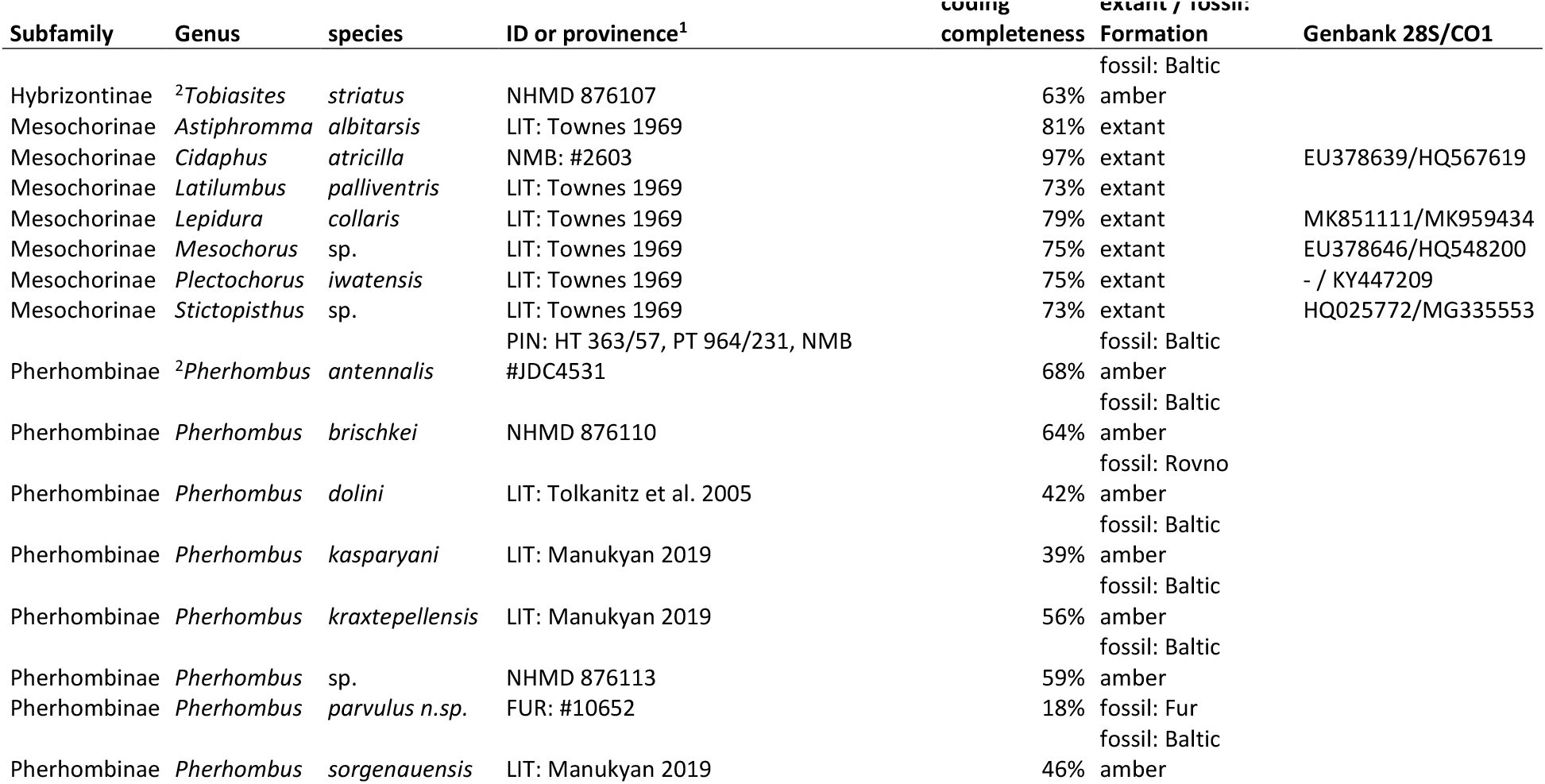
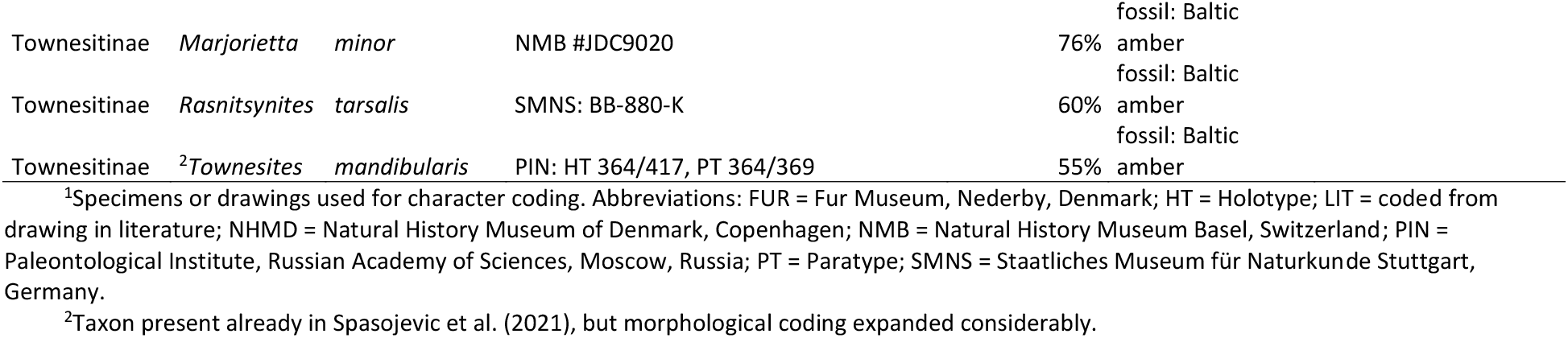
Added taxa or taxa with expanded morphological coding in comparison to the dataset from Spasojevic et al. (2021).

Of the 222 characters coded in the morphological matrix by Spasojevic et al. (2021), we excluded 12 characters that either became uninformative under our restricted taxon sampling or consisted of large amounts of missing data and could in any case not be coded for fossils. The excluded characters are the following (numbering according to Spasojevic et al. 2021): #15 (Clypeus, apical tubercule: size); #33 (Occipital notch above foramen magnum: presence/absence); #34 (Foramen magnum, flange: width); #35 (Foramen magnum: shape); #36 (Foramen magnum: location); #91 (Intercoxal carinae, position); #92 (Hind coxa, apodeme: twisting); #93 (Propodeal denticles, presence/absence); #129 (Bullae in 2m-cu: size); #200 (Tergite 8 of female, lower anterolateral corner: shape); #203 (Tergites 8 & 9 in male: fusion); #207 (Tergite 9 in female: shape).

We added two characters that are informative about Mesochorinae and Pherhombinae: “Flagellomere 1: ratio of length to width” (continuous); and “Forewing vein 1-M+1-Rs: length compared to length of r-rs” (continuous). In another two cases, we added states to existing characters in order to account for the newly included taxa: #133 (“Distal abscissa of Rs (4-Rs): shape”): (state 5) evenly arched towards 2-R1; #163 (“Tergite 1: shape from above”): (state 3) no clear separation of postpetiolus, constriction in the anterior half, thus expanding again towards the anterior margin. Sixteen characters were recorded as continuous characters and later on transformed to six-state, discrete characters in a linear fashion, as MrBayes only allows for a maximum of six states in ordered characters.

In the end, our matrix included 212 morphological characters from 12 fossil and 53 extant species and molecular data from two to nine genes (4326 bp) from the latter. The molecular data was added to stabilize the backbone of the ichneumonid subfamily tree, given that previous analyses with morphological data only resulted in poor resolution of deeper nodes in the tree (Klopfstein and Spasojevic 2019). The dataset is available as Supplementary File S2 from the Dryad Digital Repository: https://doi.org/10.5061/dryad.[NNNN] and from TreeBASE under study TB2:S28484).

### Phylogenetic analysis

A Bayesian analysis of the combined molecular and morphological partition was conducted in MrBayes 3.2 (Ronquist et al. 2012b). We used the Mkv model for the morphological partition (Lewis 2001), with 57 of the characters treated as ordered and rate-variation among characters modelled under a gamma distribution. This model was preferred over an unordered or an equal-rates model in analyses of a precursor dataset (Klopfstein and Spasojevic 2019). The molecular data was partitioned as in Spasojevic et al. (2021) and analysed under a reversible-jump MCMC substitution model (Huelsenbeck et al. 2004), including a gamma-distribution and invariant sites to model among-site rate variation.

Four independent runs of four Metropolis-coupled chains each were run for 100 million generations and convergence was assessed by inspection of the likelihood plots, effective sample sizes, potential scale-reduction factors and average standard deviation of split frequencies (ASDSF). Convergence was difficult to attain, especially on topology, with ASDSF values among the four independent runs not dropping below 0.029. This was probably due to the fossils acting as rogue taxa, a suspicion that was confirmed when comparing consensus trees with fossils included or excluded. We thus also constructed consensus trees for each of the four runs independently to make sure that our results were not influenced unduly by different runs getting stuck on different topology islands. We conservatively excluded the first half of each run as burn-in. The tree was rooted with Xoridinae as outgroup, as suggested by recent phylogenetic analyses (Bennett et al. 2019, Klopfstein et al. 2019a).

### Rogue Plots

To calculate and illustrate alternative placements of our fossil, we constructed RoguePlots (Klopfstein and Spasojevic 2019). To that end, we sampled 1000 evenly spaced trees from the post-burn-in period of all four MCMC runs in the Bayesian analysis, both separately and combined, using a custom bash script. These trees were input into the create.rogue.plot function in the rogue.plot package in R (R Core Team 2014), together with the consensus tree of all the other taxa, excluding the fossil in question.

## Results

### Morphometric Analysis

Both studied wing venation ratios clearly indicate that the new fossil species belongs to the subfamily Pherhombinae rather than Mesochorinae (Fig. 1), with both ratios allowing for a clear separation of the two subfamilies. This finding is robust to the addition of forewing information of the remaining, incomplete Pherhombinae and Mesochorinae fossils (Supplementary File S3).

**Figure 1.**
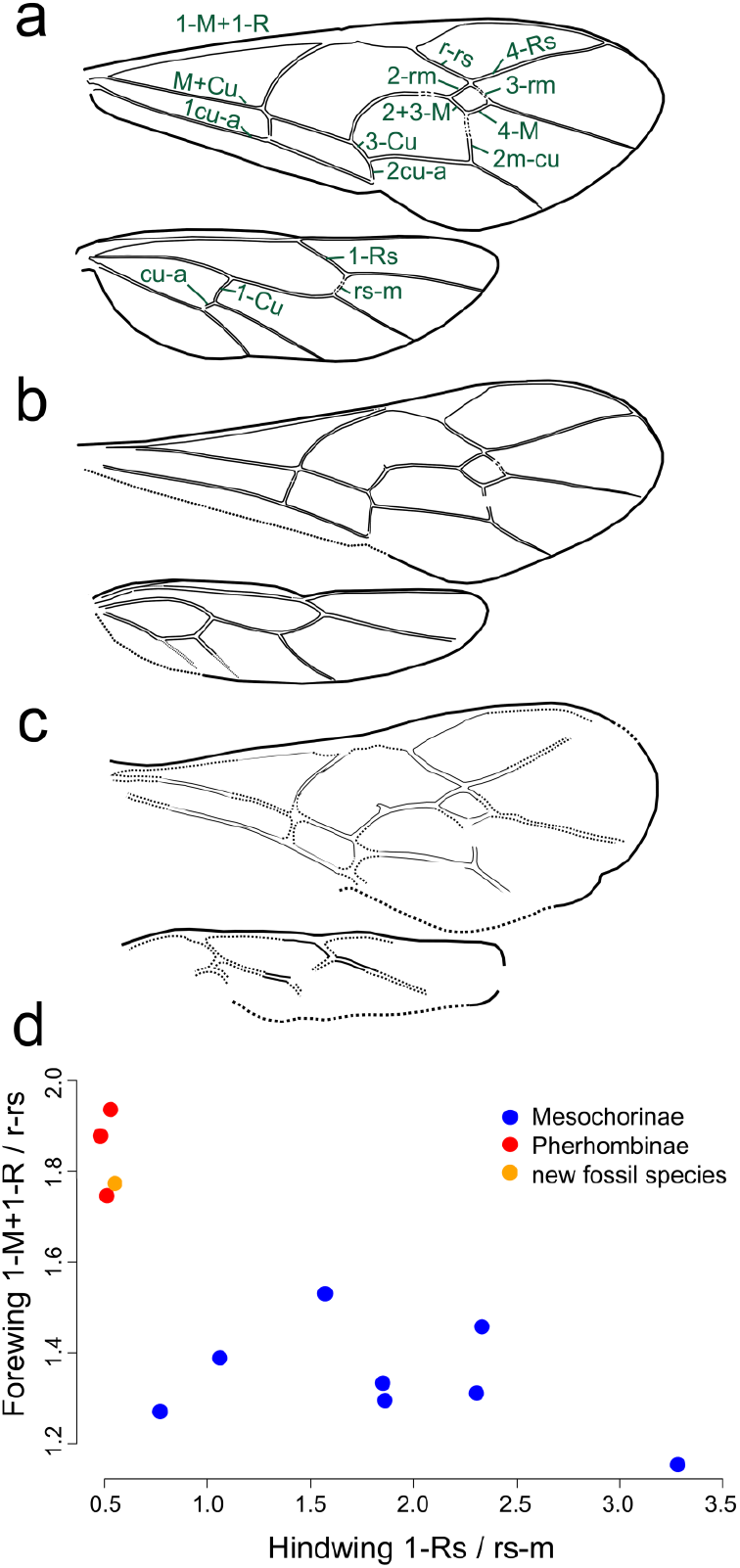
Scatterplot of two measurements ratios, forewing veins 1-M+1-Rs / r-rs and hindwing veins 1-Rs / rs-m. The new fossil species groups together with Pherhombinae. The inlaid drawings depict the respective wing vein lengths.

### Phylogeny and Rogue Plot

The phylogenetic and RoguePlot analyses undoubtedly assign the new fossil species to Pherhombinae (Fig. 2), with 1.0 posterior probability in each of the four independent MCMC runs, while there was zero support for an alternative placement with Mesochorinae. With a posterior probability of 0.64 (0.625 to 0.668 in the four runs), the new species is placed as the sister taxon to the other *Pherhombus* species. The remaining 0.36 probability is distributed to branches within the clade formed by the other *Pherhombus* species. Furthermore, the main subfamily clades (Ichneumoniformes, Pimpliformes and Ophioniformes) were all recovered in the analysis, although sometimes with low support (Fig. 2). As suggested earlier, Pherhombinae are placed in a clade with Hybrizontinae and Townesitinae within the higher Ophioniformes. Support is surprisingly high for a sister relationship between Pherhombinae and Hybrizontinae, given that the former only had morphological data (pp = 0.87). The clade in which these two group with Townesitinae is somewhat less well supported (pp = 0.62), as is their placement among the higher ophioniform subfamilies (Anomaloninae, Campopleginae, Cremastinae, Ophioninae; pp = 0.64). These results were consistent across all four independent runs.

**Figure 2.**
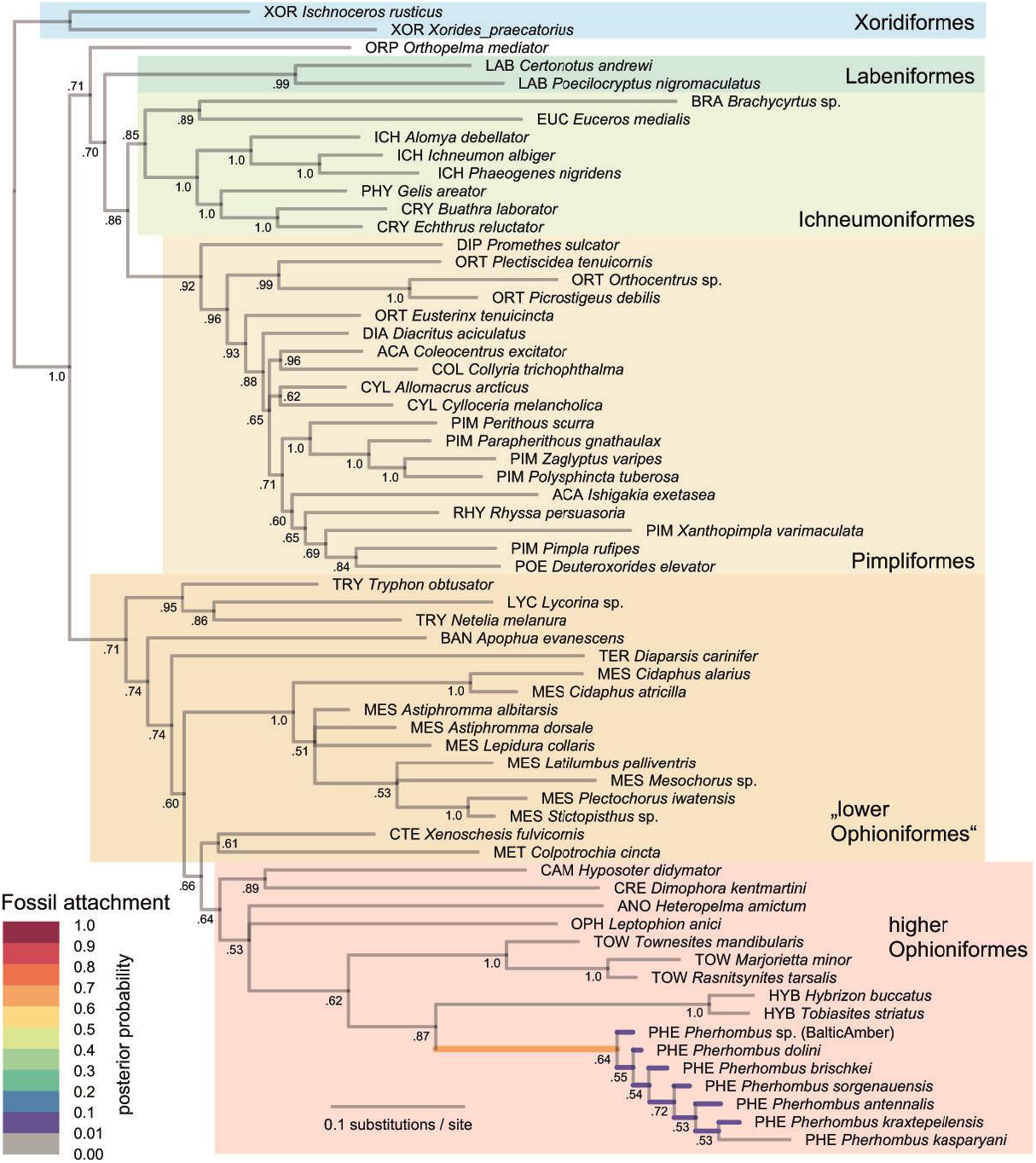
Bayesian phylogenetic analysis of combined morphological and molecular dataset, including RoguePlot showing probability of placement of *Pherhombus parvulus* n. sp. Branch colours represent posterior probabilities of attachment of the fossil to a particular branch, and support values next to nodes indicate posterior probabilities. The three-letter code in front of the taxon names denotes subfamily affiliation as follows: ACA Acaenitinae, ANO Anomaloninae, BAN Banchinae, BRA Brachycyrtinae, CAM Campopleginae, COL Collyriinae, CRE Cremastinae, CRY Cryptinae, CTE Ctenopelmatinae, CYL Cylloceriinae, DIA Diacritinae, DIP Diplazontinae, EUC Eucerotinae, HYB Hybrizontinae, ICH Ichneumoninae, LAB Labeninae, LYC Lycorininae, MES Mesochorinae, MET Metopiinae, OPH Ophioninae, ORP Orthopelmatinae, ORT Orthocentrinae, PHY Phygadeuontinae, PHE Pherhombinae, PIM Pimplinae, POE Poemeniinae, RHY Rhyssinae, TER Tersilochinae, TOW Townesitinae, TRY Tryphoninae, XOR Xoridinae.

### Systematic Palaeontology

> Family Ichneumonidae Latreille, 1802
>
> Subfamily Pherhombinae Kasparyan, 1988
>
> Genus *Pherhombus* Kasparyan, 1988
>
> *Pherhombus parvulus* **n. sp**.
>
> Figure 3

**Figure 3.**
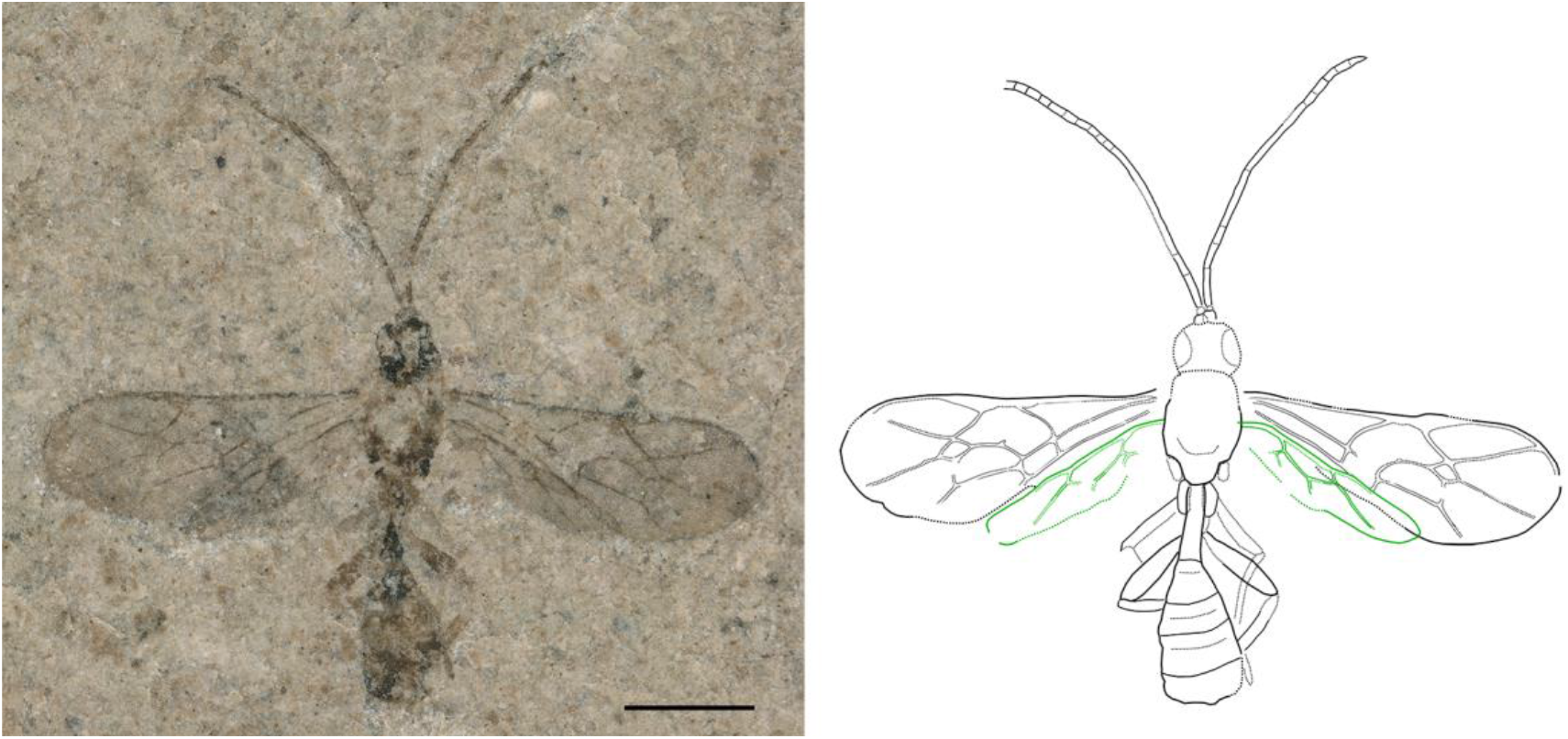
*Pherhombus parvulus* (holotype), microscope image of part 10652_A (left); interpretative drawing based on part and counterpart (right; a photograph of the counterpart is provided as Supplementary File S4). Solid lines imply a high certainty of interpretations, while dotted lines indicate interpolations or uncertain interpretations. Hindwings are shown in green to improve clarity. Differences in line width are used to visualise small structures and do not imply varying certainty. The scale bar indicates 1 mm.

#### Material

Holotype, #10652, part and counterpart; sex unknown. Part and counterpart about equally informative, often showing complementary structures (e.g. first tergite). Collector: Jan Verkleij. Deposited at Fur Museum, Nederby.

#### Type horizon and locality

The fossil was found in Denmark, Morsø Kommune, Ejerslev in cement stone which has a geological age of about 54 Ma (early Eocene).

#### Etymology

In latin, “parvulus” is the diminutive of “parvus” which means small or tiny. This refers to the fact that the fossil species is only 3.3 mm long, which is about half the size of all other described Pherhombinae.

#### Diagnosis

##### Taxonomic placement

Due to the nearly complete preservation of the forewing venation, this species can be placed within the Ichneumonidae with certainty, which are distinguished from the related Braconidae by lacking vein 1-Rs+M (sometimes with the exception of a short remain, called ramulus) and by the presence of 2m-cu. The rhombic aerolet is the probably most conspicuous character visible in this fossil; it is only shared by members of the subfamilies Mesochorinae and Pherhombinae. Several characters, such as the low number of antennal segments, the forewing 1-M+1-Rs to r-rs ratio, the hindwing 1-Rs to rs-m ratio and the elongated and parallel-sided first tergite give evidence for the placement within the monotypic Pherhombinae. Even though the new species was placed with the highest probability as a stem representative, there is currently not sufficient morphological evidence that this species should be placed within a new genus. This placement is supported both by the morphometric and Bayesian phylogenetic analyses (Figs 1 and 2).

##### Species diagnosis

This species is very similar both in wing venation and shape of the first tergite to many species of *Pherhombus*. Nevertheless, it can be easily distinguished from all currently described species by its small size (body length: 3.3 mm, forewing length: 2.9 mm), with all other *Pherhombus* speices ranging in body length from 6.7 to more than 9 mm and forewing length from 4.7 to 9.5 mm (Tolkanitz et al. 2005, Manukyan 2019). It is further distinguished from most other species by the parallel-sided antennal segments, which are otherwise widened towards the apex in all species except perhaps *P. kraxtepellensis* and *P. kasparyani* (Manukyan, 2019), whose antennae are widest around the 8^th^ or 9^th^ flagellomere and only slightly expanded apically. The new species can be distinguished from these two by the presence of a distinct ramulus and by the hyaline wings (smoky in *P. kasparyani*).

#### Description

##### Preservation

Dorsal view. Head only partially preserved, antennae nearly complete with partly clear segmentation. Mesosoma not well preserved, hardly any characters visible except possible hind border of mesoscutum; wings stretched out flat, all four wings nearly complete; partial mid and hind legs visible. Metasoma anteriorly almost complete but segmentation posteriorly unclear; posterior part of metasoma ending abruptly or incomplete, genitalia not visible.

Body 3.3 mm; fossil in different shades of brown; strongly sclerotized parts, such as head or first tergite, distinctly darker than rest, wings hyaline.

*Head* deformed, no detailed structures distinguishable. Antenna slender, with about 20 flagellomeres (+/- 3); scape and pedicel of normal dimensions (as far as visible), first flagellomere almost 7.0 times longer than apically wide.

*Mesosoma* rather short and stout; triangular dark patches at forewing base probably corresponding to axial sclerites. Forewing 2.9 mm; areolet closed, rhombic, 3r-m with a bulla at posterior end; 4-Rs straight; ramulus present, slightly longer than width of surrounding veins; pterostigma 4.5 x longer than wide; radial cell 3 x longer than wide; 1cu-a meeting M+Cu opposite 1-M&1-RS; 2m-cu nearly straight, somewhat inclivous, probably with a single large bulla in anterior third or half; 3-Cu about 0.75 x as long as 2cu-a. Hindwing 1-Rs 0.47 x as long as rs-m; 2-Rs tubular on entire length (not counting last 10%); 1-Cu clearly shorter than cu-a. Mid leg very slender, both coxa, femur and parts of tibia and tarsus visible. Hind leg with very long coxa, at least 2.1 x longer than wide; both femur and parts of tibia preserved, femur rather elongated, more than 5.0 x longer than wide.

*Metasoma* appearing somewhat club-shaped, with widest part close to posterior end; tergite 1 slightly more than 4 x longer than wide, narrow and parallel sided; tergite 2 transverse, 0.75 x as long as wide. Posterior metasomal segments appear truncated, lack of ovipositor suggests a male, but incomplete preservation also possible.

## Discussion

### Integrative analysis facilitates firm fossil placement

The integrative approach we followed here facilitated a firm placement of the newly described fossil species in an extinct subfamily. The discussion whether to work with molecular or morphological data is omnipresent in entomological systematics. In most cases, integrative approaches are beneficial for taxonomic studies as they manage to grasp more of the available information (Schlick-Steiner et al. 2010, Yeates et al. 2011, Wang et al. 2015, Gokhman 2018). This is just as true for the phylogenetic placement of fossils – complementing morphological analyses with molecular data for recent taxa gives higher stability especially to the backbone of phylogenetic trees (Nylander et al. 2004, Ronquist et al. 2012a, Spasojevic et al. 2021). However, placement of fossils is only possible if extensive morphological data is included for extant taxa as well, because extant taxa with missing morphological information can attract fossils in phylogenetic analyses (Spasojevic et al. 2021). Thus, morphological data should be coded for all included extant taxa, which proved to be a powerful approach in our study.

A potential drawback of Bayesian phylogenetic inference is the difficulty to directly assess the impact of individual characters on the outcome. To make sure that an analysis was not biased by few characters, several steps are possible. Character state changes can be mapped onto a phylogeny, characters can be excluded in the analysis in order to check for their impact, or additional analyses based on only a few characters can be conducted to assess their respective signal. In our case, we separately analysed morphometric data on wing venation of Pherhombinae and Mesochorinae and thus identified two ratios with high information content with regard to the differentiation of these two groups. With very few exceptions, wing venation characters have not received much attention in subfamily identification in Ichneumonidae (Broad et al. 2018, but see Li et al. 2019), and our results suggest that they should be explored in more detail in the future, maybe even in the framework of geometric morphometrics.

Not only an overrated single character, but also a systematic bias could alter the result of the analysis. In the study at hand, a potential source of systematic bias is body size, which often influences several traits at once (Minelli and Fusco 2019). Miniaturisation effects often include parallel character loss, morphological simplification, and allometric effects that might change morphometric ratios (Gould 1966, Klopfstein et al. 2015, Knauthe et al. 2016). Therefore, *P. parvulus* could have been placed in the clade of Pherhombinae, Townesitinae and Hybrizontinae because all of them are rather small ichneumonids. Indeed, some of the character states that support this placement might be related to a reduction in size, for example the reduced number of palpal segments and reduced mandibles. However, other characters are less likely to be a result of miniaturization, such as the elongate hind coxae or shape of first tergite (for more characters, see next section). Also, we found that the wing vein ratios of the new species are very similar to those of the other *Pherhombus* species, even though these are distinctly larger in body size. Furthermore, other subfamilies with many small-bodied taxa (e.g. Orthocentrinae, Phygadeuontinae, Campopleginae) did not attract the new fossil species at all (Fig. 2). We thus consider the placement as reliable, even though further research is needed to support the close relationship between the three subfamilies.

Another possible source of systematic bias is the heterogeneous origin of morphological data used in this study. Some extant taxa were studied directly and thus with detailed morphological data, some were coded based only on drawings and descriptive texts (Townes 1971). Most amber fossils that we studied were well preserved and thus show rather complete coding, often approaching extant taxa with respect to completeness (Table 1); but only a few characters could be coded for the new rock fossil species. Previous studies using the total-evidence dating framework and thus working with similarly incomplete data matrices found that even poorly coded fossils can contribute considerably to an analysis, while missing data did not seem to negatively affect the outcome (Ronquist et al. 2012a). Similarly, we here found no evidence that heterogeneous completeness biased the phylogenetic analysis, since even subfamilies which included species from very different data sources were retrieved with high support, and the new rock fossil species was placed very confidently with the amber Pherhombinae in the phylogenetic tree.

Resolution of the phylogeny reconstructed here is rather high, considering that inference of considerable portions of the tree were only informed by morphology, especially in the extinct Pherhombinae and Townesitinae. The analysis of morphological characters in a phylogenetic context of course always relies to a certain extent on the availability of an appropriate model of character evolution (Lewis 2001, Klopfstein et al. 2015, Wright et al. 2016). However, the high congruence between the morphological and molecular partition in our dataset suggests that model mismatch is not strongly misleading our results (Klopfstein and Spasojevic 2019), although our analysis would certainly profit from the development of more refined models of morphological evolution.

### Phylogenetic support for Kasparyan’s hypothesis

Our phylogenetic analysis supports Kasparyans’s hypothesis (1988, Kasparyan 1994) that Pherhombinae, Townesitinae and Hybrizontinae form a clade, which in our analysis was located within the higher Ophioniformes. The three subfamilies share several derived character states from different parts of the body, though most of them are not entirely exclusive to the three. The strongly convex clypeus also occurs in Orthocentrinae, an extant subfamily that consists exclusively of small-bodied taxa. However, it is less convex in Orthocentrinae and much more narrow in Hybrizontinae than in the other subfamilies. The maxillar and labial palps have a reduced number of segments in Hybrizontinae, Pherhombinae and in one of the two tribes of Townesitinae, but not in Orthocentrinae. Vein r-rs in the forewing is conspicuously shortened in all three subfamilies, as is 1RS in the hindwing in those species where it is visible (Fig. 1). The hind coxa is rather elongate, and the first tergite is narrow and elongate in all three subfamilies, although more strongly so in Pherhombinae.

In our tree, Hybrizontinae and Pherhombinae group together, which was also Kasparyan’s initial suggestion (1988). Later, he apparently changed his mind, assuming that Pherhombinae and Townesitinae were more closely related (Kasparyan 1994, Manukyan 2019), which was also the outcome of a previous phylogenetic study with a more sparse taxon sampling (Spasojevic et al. 2021). Our current analysis now once more revives the initial suggestion of close ties between Pherhombinae and the highly derived Hybrizontinae, and several character states support this relationship. The mandibles are reduced to a flap-like structure in Hybrizontinae, Pherhombinae and in some species of *Orthocentrus*, while being complete with two teeth in Townesitinae. The metasomal cavity is moved upwards with respect to the metacoxal cavities in both Pherhombinae and Hybrizontinae, but not Townesitinae, a state that is otherwise only present in Labeninae. Finally, the hind coxa is much more strongly elongate in Pherhombinae and Hybrizontinae than in Townesitinae.

Based on Kasparyan’s (1994) suggested relationships, Manukyan (2019) inferred that the three subfamilies have split in the mid to late Eocene. Considering that the new fossil species is clearly placed after the splitting of Pherhombinae and Hybrizontinae and given the age of the Fur Formation, we can provide good evidence that the last common ancestor of the three groups lived much earlier than that, probably already in the Palaeocene or earlier. This suggestion is supported by the outcome of a recent dating analysis, which found that most subfamilies of Darwin wasps already diversified in the Mesozoic (Spasojevic et al. 2021). Studies of late Cretaceous ambers should thus consider the possibility that members of these subfamilies or stem-lineages there-of already occurred before the K-Pg mass extinction.

### Presence of Pherhombinae in rock deposit from earliest Eocene

Defining the age span of the monotypic Pherhombinae is rather difficult because up to now, only amber fossil species were known. *Pherhombus dolini* from Rovno amber (Tolkanitz et al. 2005) anchors the upper age limit of the subfamily in the upper Eocene (33.9–37.8 Ma). The other *Pherhombus* species are known only from Baltic amber and thus do not provide much information about the age span of Pherhombinae, as the age of Baltic amber is still highly controversial in the paleontological community (Standke 1998, Schulz 1999, Weitschat and Wichard 2010, Sadowski et al. 2017, Tolkanitz and Perkovsky 2018). Our finding of *P. parvulus* from a lowermost Eocene rock deposit pushes back the lower age maximum of *Pherhombus* to about 54 Ma, thus leading to a minimal age span of the subfamily of nearly 20 million years. Even though this is a long time period, it should not be considered as unlikely, given that other ichneumonid genera existed for much longer periods, e.g., *Phaenolobus* or *Xanthopimpla* (56 and 54 Ma–present: Piton 1940, Klopfstein in press). Considering these long time windows for ichneumonid genera, making inferences from *P. parvulus* on the age estimate of Baltic amber appears unwarranted; however, its finding at least allows for the possibility that Baltic amber might be considerably older than Rovno amber. On the other hand, our phylogenetic analysis suggests that *P. parvulus* is the sister taxon to all other *Pherhombus* species, which is congruent with the notion that it lived much earlier than its congeners. Interestingly, our new species is not just the oldest but also the smallest species known of this subfamily, which is remarkable given that it was found in rock rather than amber, even though the latter is typically known for a bias towards small-bodied taxa. This might indicate that Pherhombinae increased their body size over time, even though this conclusion is somewhat shaky given the low number of known species.

The new Pherhombinae fossil described here exemplifies just how poorly studied ichneumonid fossils still are. Future studies might reveal an even more extensive temporal distribution of this enigmatic subfamily and might provide further clues as to their ecology and evolution. Furthermore, each described and properly placed fossil Darwin wasp can contribute to the proper calibration of the phylogenetic tree of this hyperdiverse insect group and thereby improve our knowledge of its evolution and diversification.

## Acknowledgements

We are grateful to René Sylvestersen of the Fur Museum in Nederby, Denmark, for providing the fossil described here. We thank Lars Vilhelmsen (Natural History Museum of Denmark, Copenhagen), Dmitry Kopylov (Paleontological Institute, Russian Academy of Sciences, Moscow, Russia) and Michael Rasser (Staatliches Museum für Naturkunde Stuttgart, Germany) for access to amber fossils from their collections for comparison. Tamara Spasojevic and Bastien Mennecart provided valuable feedback on an earlier version of the manuscript.

Computations were performed on the HPC cluster UBELIX of the University of Bern, Switzerland (http://www.id.unibe.ch/hpc).

## Data availability

All data used in this study is available from the Dryad Digital Repository: https://doi.org/10.5061/dryad.[NNNN]

## References

Ansorge J. 1993. Schlupfwespen aus dem Moler Dänemarks – ein aktualistischer Vergleich. Fossilien. Korb, Goldschneck-Verlag, p. 111–113.

Belshaw R, Grafen A, Quicke DLJ. 2003. Inferring life history from ovipositor morphology in parasitoid wasps using phylogenetic regression and discriminant analysis. Zool J Linn Soc, 139:213–228 doi: https://doi.org/10.1046/j.1096-3642.2003.00078.x.

Bennett AMR, Cardinal S, Gauld ID, Wahl DB. 2019. Phylogeny of the subfamilies of Ichneumonidae (Hymenoptera). J Hym Res, 71:1–156 doi: https://doi.org/10.3897/jhr.71.32375.

Broad GR, Shaw MR, Fitton MG. 2018. Ichneumonid wasps (Hymenoptera: Ichneumonidae): their classification and biology. Handbooks for the identification of British insects, 7:1–418.

Brues CT. 1910. The parasitic Hymenoptera of the Teritary of Florissant, Colorado. Bulletin of the Museum of Comparative Zoology at Harvard University, 54:1–125.

Brues CT. 1923. Some new fossil parasitic Hymenoptera from Baltic amber. Proceedings of the American Academy of Arts and Sciences, 58:327–346 doi: https://doi.org/10.2307/20025999.

Bukejs A, Alekseev VI, Pollock DD. 2019. Waidelotinae, a new subfamily of Pyrochroidae (Coleoptera: Tenebrionoidea) from Baltic amber of the Sambian peninsula and the interpretation of Sambian amber stratigraphy, age and location. Zootaxa, 4664:261–273 doi: https://doi.org/10.11646/zootaxa.4664.2.8.

Chambers LM, Pringle M, Fitton G, Larsen LM, Pedersen AK, Parrish R. 2003. Recalibration of the Palaeocene-Eocene boundary (PE) using high precision U-Pb and Ar-Ar isotopic dating. EGS-AGU-EUG Joint Assembly. Nice.

Dunlop JA, Vlaskin AP, Marusik Y. 2019. Comparing arachnids in Rovno amber with the Baltic an Bitterfeld deposits. Paleontological Journal, 53:92–101 doi: https://doi.org/10.1134/S0031030119100034.

Forbes AA, Bagley RK, Beer MA, Hippee AC, Widmayer HA. 2018. Quantifying the unquantifiable: why Hymenoptera – not Coleoptera – is the most speciose animal order. BMC Ecology, 18:21 doi: https://doi.org/10.1186/s12898-018-0176-x.

Gokhman VE. 2018. Integrative taxonomy and its implications for species-level systematics of parasitoid Hymenoptera. Entomological Review, 98:834–864 doi: https://doi.org/10.1134/S0013873818070059.

Gould SJ. 1966. Allometry and size in ontogeny and phylogeny. Biol Rev, 41:587–640 doi: https://doi.org/10.1111/j.1469-185X.1966.tb01624.x.

Henriksen KL. 1922. Eocene insects from Denmark. Danmarks geologiske Undersøgelse, 2:1–36 doi: https://doi.org/10.34194/raekke2.v37.6823.

Huelsenbeck JP, Larget B, Alfaro ME. 2004. Bayesian phylogenetic model selection using reversible-jump Markov chain Monte Carlo. Mol Biol Evol, 21:1123–1133 doi: https://doi.org/10.1093/molbev/msh123.

Kasparyan DR. 1988. A new subfamily and two new genera of ichneumonids (Hymenoptera, Ichneumonidae) from Baltic amber. [In Russian]. Proceedings of the Zoological Institute Leningrad, 175:38–42.

Kasparyan DR. 1994. A review of the Ichneumon flies of Townesitinae subfam. nov. (Hymenoptera, Ichneumonidae) from the Baltic amber. Paleontological Journal, 28:114–126.

Klopfstein S. in press. High diversity of pimpline parasitoid wasps (Hymenoptera, Ichneumonidae, Pimplinae) in the Fur Formation in Denmark. Geodiversitas.

Klopfstein S, Langille B, Spasojevic T, Broad GR, Cooper SJB, Austin AD, Niehuis O. 2019a. Hybrid capture data unravels a rapid radiation of pimpliform parasitoid wasps (Hymenoptera: Ichneumonidae: Pimpliformes). Syst Entomol, 44:361–383 doi: https://doi.org/10.1111/syen.12333.

Klopfstein S, Santos B, Shaw MR, Alvarado M, Bennett AMR, Dal Pos D, Giannotta M, Herrera Florez AF, Karlsson D, Khalaim AI, et al. 2019b. Darwin wasps: a new name heralds renewed efforts to unravel the evolutionary history of Ichneumonidae. Entomological Communications, 1:ec01006.doi: https://doi.org/10.37486/2675-1305.ec01006.

Klopfstein S, Spasojevic T. 2019. Illustrating phylogenetic placement of fossils using RoguePlots: An example from ichneumonid parasitoid wasps (Hymenoptera, Ichneumonidae) and an extensive morphological matrix. PLoS One, 14:e0212942.doi: https://doi.org/10.1371/journal.pone.0212942.

Klopfstein S, Vilhelmsen L, Ronquist F. 2015. A nonstationary Markov model detects directional evolution in hymenopteran morphology. Syst Biol, 64:1089–1103 doi: https://doi.org/10.1093/sysbio/syv052.

Knauthe P, Beutel RG, Hörnschemeyer T, Pohl H. 2016. Serial block-face scanning electron microscopy sheds new light on the head anatomy of an extremely miniaturized insect larva (Insecta, Strepsiptera). Arthropod Syst Phylogeny, 74:107–126.

Larsson SG. 1975. Palaeobiology and mode of burial of the insects of the lower Eocene Mo-clay of Denmark. Bulletin fo the geological society of Denmark, 24:193–209.

Lewis PO. 2001. A likelihood approach to estimating phylogeny from discrete morphological character data. Syst Biol, 50:913–925 doi: https://doi.org/10.1080/106351501753462876.

Li L, Shih PJM, Kopylov DS, Li D, Ren D. 2019. Geometric morphometric analysis of Ichneumonidae (Hymenotpera: Apocrita) with two new Mesozoic taxa from Myanmar and China. Journal of Systematic Palaeontology, DOI: 10.1080/14772019.2019.1697903 doi: https://doi.org/10.1080/14772019.2019.1697903.

Manukyan AR. 2019. New data on ichneumon wasps of the subfamily Pherhombinae (Hymenoptera, Ichneumonidae) from Baltic amber, with descriptions of three new species. Entomological Review, 99:1324–1338 doi: https://doi.org/10.1134/S0013873819090112.

McIntosh WC, Geissmann JW, Chapin CE, Kunk MJ, Henry CD. 1992. Calibration of the latest Eocene-Oligocene geomagnetic polarity time scale using 40Ar/39Ar dated ignimbrites. Geology, 20:459–463 doi: https://doi.org/10.1130/0091-7613(1992)020<0459:COTLEO>2.3.CO;2.

Minelli A, Fusco G. 2019. No limits: Breaking constraints in insect minituarization. Arthropod Struct Dev, 48:4–11 doi: https://doi.org/10.1016/j.asd.2018.11.009.

Nylander JAA, Ronquist F, Huelsenbeck JP, Nieves-Aldrey JL. 2004. Bayesian phylogenetic analysis of combined data. Syst Biol, 53:47–67 doi: https://doi.org/10.1080/10635150490264699.

Perkovsky EE, Rasnitsyn AP, Vlaskin AP, Taraschuk MV. 2007. A comparative analysis of the Baltic and Rovno amber arthropod faunas: representative samples. African Invertebrates, 48:229–245 doi: 10.10520/EJC84578.

Piton L. 1940. Paléontologie du gisement éocène de Menat (Puy-de-Dôme) (Flore et Faune). Faculté des Sciences. Clermont-Ferrand, Université de Clermont p. 303.

Pyron RA. 2011. Divergence time estimation using fossils as terminal taxa and the origins of Lissamphibia. Syst Biol, 60:466–481.

R Core Team. 2014. R: A language and environment for statistical computing. Vienna, Austria, R Foundation for Statistical Computing.

Ritzkowski S. 1997. K-Ar-Altersbestimmungen der bernsteinführenden Sedimente des Samlandes (Paläogen, Bezirk Kaliningrad). Metalla, Bochum, 66:19–23.

Ronquist F, Forshage M, Häggqvist S, Karlsson D, Hovmöller R, Bergsten J, Holston K, Britton T, Abenius J, Andersson B, et al. 2020. Completing Linnaeus’s inventory of the Swedish insect fauna: Only 5,000 species left? PLoS One, 15:e0228561.doi: https://doi.org/10.1371/journal.pone.0228561.

Ronquist F, Klopfstein S, Vilhelmsen L, Schulmeister S, Murray DL, Rasnitsyn AP. 2012a. A total-evidence approach to dating with fossils, applied to the early radiation of the Hymenoptera. Syst Biol, 61:973–999 doi: http://dx.doi.org/10.1093/sysbio/sys058.

Ronquist F, Teslenko M, Van der Mark P, Ayres DL, Darling A, Höhna S, Larget B, Liu L, Suchard MA, Huelsenbeck JP. 2012b. MrBayes 3.2: Efficient Bayesian phylogenetic inference and model choice across a large model space. Syst Biol, 61:539–542 doi: https://doi.org/10.1093/sysbio/sys029.

Rust J. 1998. Biostratinomie von Insekten aus der Fur-Formation von Dänemark (Moler, oberes Paleozän / unteres Eozän). Paläontologische Zeitschrift, 72:41–58 doi: https://doi.org/10.1007/BF02987814.

Rust J. 2000. Fossil record of mass moth migration. Nature, 405:530–531 doi: https://doi.org/10.1038/35014733.

Sadowski EM, Schmidt AR, Saeyfullah LJ, Kunzmann L. 2017. Conifers of the ‘Baltic amber forest’ and their palaeoecological significance. Stapfia, 106:2–73.

Schlick-Steiner BC, Steiner FM, Seifert B, Stauffer C, Christian E, Crozier RH. 2010. Integrative taxonomy: a multisource approach to exploring biodiversity. Annu Rev Entomol, 55:421–438 doi: https://doi.org/10.1146/annurev-ento-112408-085432.

Schulz W. 1999. The Baltic Amber in Qaternary sediments, a review of its occurrence, the largest specimens and amber museums. Archiv für Geschiebekunde, 2:459–478.

Spasojevic T, Broad GR, Bennett AMR, Klopfstein S. 2018. Ichneumonid parasitoid wasps from the Early Eocene Green River Formation: five new species and a revision of the known fauna (Hymenoptera, Ichneumonidae). Paläontologische Zeitschrift, 92:35–63 doi: https://doi.org/10.1007/s12542-017-0365-5.

Spasojevic T, Broad GR, Sääksjärvi IE, Schwarz M, Ito M, Matsumoto R, Hopkins T, Klopfstein S. 2021. Mind the outgroup and bare branches in total-evidence dating: a case study of pimpliform Darwin wasps (Hymenoptera, Ichneumonidae). Syst Biol, 70:322–339 doi: https://doi.org/10.1093/sysbio/syaa079.

Standke G. 1998. Die Tertiärprofile der Samländischen Bernsteinküste bei Rauschen. Schriftenreihe für Geowissenschaften, 7:93–133.

Tolkanitz VI, Narolsky NB, Perkovsky EE. 2005. A new species of parasite wasp of the genus Pherhombus (Hymenoptera, Ichneumonidae, Pherhombinae) from the Rovno Amber [in Russian]. Paleontologicheskii Zhurnal, 5:50–52.

Tolkanitz VI, Perkovsky EE. 2018. First record of the late Eocene Ichneumon fly Rasnitsynites tarsalis Kasparyan (Ichneumonidae, Townesitinae) in Ukraine confirms correlation of the upper Eocene Lagerstätten. Paleontological Journal, 52:31–34 doi: https://doi.org/10.1134/S0031030118010136.

Townes HK. 1971. The genera of Ichneumonidae, Part 4. Memoirs of the American Entomological Institute, 17:1–372.

Wang Y, Nansen C, Zhang Y. 2015. Integrative insect taxonomy based on morphology, mitochondrial DNA, and hyperspectral reflectance profiling. Zool J Linn Soc, 177:378–394 doi: https://doi.org/10.1111/zoj.12367.

Weitschat W, Wichard W. 2010. Baltic Amber. In: Penney D editor. Biodiversity of fossils in amber from the major world deposits. Manchester, Siri Scientific Press, p. 80–115.

Westerhold T, Röhl U, McCarren HK, Zachos JC. 2009. Latest on the absolute age of the Paleocene™Eocene Thermal Maximum (PETM): New insights from exact stratigraphic position of the key ash layers +19 and ™17. Earth and Planetary Science Letters, 287:412–419 doi: https://doi.org/10.1016/j.epsl.2009.08.027.

Wright AM, Lloyd GT, Hillis DM. 2016. Modeling character change heterogeneity in phylogenetic analyses of morphology through the use of priors. Syst Biol, 65:602–611 doi: https://doi.org/10.1093/sysbio/syv122.

Yeates DK, Seago A, Nelson L, Cameron SL, Joseph L, Trueman JWH. 2011. Integrative taxonomy, or iterative taxonomy? Syst Entomol, 36:209–217.

Yu DS, Van Achterberg C, Horstmann K. 2016. Taxapad 2016. Ichneumonoidea 2015 (Biological and taxonomical information), Taxapad Interactive Catalogue Database on flash-drive. Nepean, Ottawa, Canada, http://www.taxapad.com.

